# Causal inference in cancer epidemiology: what is the role of Mendelian randomization?

**DOI:** 10.1101/223966

**Authors:** James Yarmolinsky, Kaitlin H Wade, Rebecca C Richmond, Ryan J Langdon, Caroline J Bull, Kate M Tilling, Caroline L Relton, George Davey Smith, Richard M Martin

**Affiliations:** Integrative Epidemiology Unit, University of Bristol, Bristol, UK; Population Health Sciences, Bristol Medical School, University of Bristol, Bristol, UK

## Abstract

Observational epidemiological studies are prone to confounding, measurement error, and reverse causation, undermining their ability to generate reliable causal estimates of the effect of risk factors to inform cancer prevention and treatment strategies. Mendelian randomization (MR) is an analytical approach that uses genetic variants to proxy potentially modifiable exposures (e.g. environmental factors, biological traits, and druggable pathways) to permit robust causal inference of the effects of these exposures on diseases and their outcomes. MR has seen widespread adoption within population health research in cardio-metabolic disease, but also holds much promise for identifying possible interventions (e.g., dietary, behavioural, or pharmacological) for cancer prevention and treatment. However, some methodological and conceptual challenges in the implementation of MR are particularly pertinent when applying this method to cancer aetiology and prognosis, including reverse causation arising from disease latency and selection bias in studies of cancer progression. These issues must be carefully considered to ensure appropriate design, analysis, and interpretation of such studies.

In this review, we provide an overview of the key principles and assumptions of MR focusing on applications of this method to the study of cancer aetiology and prognosis. We summarize recent studies in the cancer literature that have adopted a MR framework to highlight strengths of this approach compared to conventional epidemiological studies. Lastly, limitations of MR and recent methodological developments to address them are discussed, along with the translational opportunities they present to inform public health and clinical interventions in cancer.

## Introduction

Obtaining reliable evidence of causal relationships from observational epidemiological studies remains a pervasive challenge ^1–3^. While observational studies have made fundamental contributions to understanding the primary environmental causes of various cancers (e.g., smoking and lung cancer, hepatitis B and liver cancer, asbestos and mesothelioma) ^4–6^, recent decades have seen numerous instances of apparently robust observational associations being subsequently contradicted by large chemoprevention trials ^8–16^. Notable translational failures include the ineffectiveness of beta-carotene supplementation to prevent lung cancer among smokers in the Alpha-Tocopherol, Beta-Carotene Cancer Prevention Study and vitamin E supplementation to prevent prostate cancer in the Selenium and Vitamin E Cancer Prevention Trial. Contrary to expectations from observational data, findings from both trials suggested that supplementation may increase rather than reduce the incidence of cancer ^9,17^

Part of the difficulty in translating observational findings into effective cancer prevention and treatment strategies lies in the susceptibility of conventional observational designs to various biases, such as residual confounding (due to unmeasured or imprecisely measured confounders) and reverse causation ^18,19^ These biases frequently persist despite energetic statistical and methodological efforts to address them ^20–22^, making it difficult for observational studies to reliably conclude that a risk factor is causal, and thus a potentially effective intervention target. This issue is likely further compounded by the modern epidemiological pursuit of risk factors that confer increasingly modest effects on disease risk, which can contribute to a ubiquity of spurious findings in the literature ^23–25^.

Despite these challenges, observational studies remain crucial for informing cancer prevention and treatment policy given issues in translating basic science to human populations and because intervention trials are expensive, time-consuming, and often unfeasible in a primary prevention setting. The development of novel analytical tools that can help address some of the limitations of conventional observational studies therefore remains an important field of research. One such approach known as Mendelian randomization (MR) which uses genetic variants to proxy potentially modifiable exposures has seen increased adoption within population health research and offers much promise to generate a more reliable evidence-base for cancer prevention and treatment.

### What is Mendelian randomization?

MR uses germline genetic variants as instruments (i.e., proxies) for exposures (e.g., environmental factors, biological traits, or druggable pathways) to examine the causal effects of these exposures on health outcomes (e.g., disease incidence or progression) ^26–32^ The use of genetic variants as proxies exploits their random allocation at conception (Mendel’s first law of inheritance) and the independent assortment of parental variants at meiosis (Mendel’s second law of inheritance). These natural randomization processes mean that, at a population level, genetic variants that are associated with levels of a specific modifiable exposure will generally be independent of other traits and behavioural or lifestyle factors, although several caveats exist (see **Table 1**). Analyses using genetic variants as instruments to examine associations with outcomes have a number of advantages: i) effect estimates should be less prone to the confounding that typically distorts conventional observational associations ^33^, ii) because germline genetic variants are fixed at conception, they cannot be modified by subsequent factors, thus overcoming possible issues of reverse causation, and iii) measurement error in genetic studies is often low as modern genotyping technologies provide relatively precise measurement of genetic variants, unlike the substantial (and at times differential) exposure measurement error which can accompany observational studies (e.g., due to self-report).

**Table 1.** Limitations of Mendelian randomization and techniques available to address them

| Limitation | Description | Techniques to Address Limitation |
| --- | --- | --- |
| **Limitations to robust causal inference** |  |  |
| Horizontal pleiotropy | A genetic variant affecting an outcome via a biological pathway independent of the exposure under investigation, violating the “exclusion restriction criterion” | Assessment of heterogeneity across individual SNP estimates<br>MR-Egger regression and intercept test<br>Median-based approaches<br>Mode-based approaches<br>Sensitivity analysis removing potentially pleiotropic SNPs<br>Restrict risk score to SNPs in well-characterized genes<br>Stratification by exposure status (e.g., *ALDH2* and self-reported alcohol intake) |
| Linkage disequilibrium | Linkage disequilibrium (LD) is the non-random association of alleles at different loci that are close in proximity on a chromosome. If a certain SNP is being used as an instrument for an exposure in a MR analysis, and this SNP is in LD with another SNP that affects the outcome via an independent pathway, then the assumptions for MR will be violated | LD pruning of SNPs prior to MR analysis<br>Weighted generalized linear regression<br>Perform studies in populations with different LD structures |
| Population stratification | Allele frequencies vary among populations of different genetic ancestry, and similarly, disease risk often varies among populations of different genetic ancestry, which could introduce genetic confounding into a MR analysis, potentially resulting in spurious causal estimates | Restricting analyses to individuals of a homogenous genetic ancestry<br>Genomic inflation factor calculation<br>Adjusting MR analysis by genetic ancestry or ancestry-informative principal components |
| Trait heterogeneity | For a given trait (e.g., adiposity), SNPs may influence various dimensions of this trait (e.g., both overall and visceral adiposity) but GWAS have only examined associations with a subset of these dimensions (e.g., solely BMI). This may produce misleading inferences if the aim of an analysis is to ascertain the causal effect of a particular dimension of a trait. | Better understanding of complex phenotypes<br>Multivariable MR |
| **Limitations that complicate interpretation** |  |  |
| Canalization | Developmental compensation against the effect of a genetic variant being used as an instrument that could attenuate the magnitude of an observed MR association towards the null | Knowledge of the period of life when the influence of a genetic variant(s) on an exposure may emerge can help guide whether developmental compensatory processes are plausible. For example, behavioural exposures that typically occur after fetal development (e.g., alcohol, smoking) will be unlikely to be influenced by canalization whereas in utero exposure may. There are currently no approaches for evaluating suspected canalization in MR analyses. |
| Complexity of association | Misinterpretation of MR results can arise from limited biological understanding of genetic variants utilised as IVs. Examples include interpretation of the effect of the heterozygous *ALDH2* genotype on oesophageal cancer risk (discussed in “Illustrative examples”) and previous MR analyses that have examined the effects of interleukin-6<sup>43</sup> and extracellular superoxide dismutase<sup>183</sup> on CHD risk (discussed in more detail elsewhere<sup>50</sup>). | Improved biological understanding of genetic variants with functional annotation, pathway analysis, and gene set enrichment |
| Dynastic effects | In certain circumstances, it is possible that parental genotype can confound an association of offspring genotype with offspring disease risk. For example, genetic variants influencing parental height will not only influence offspring height genotype but could also influence offspring disease risk via an independent effect of maternal height-raising alleles on the in utero environment of the offspring<sup>184,185</sup>. | Between-sibling MR design<br>Within-family MR design |
| Critical period effects | If a biomarker primarily influences disease risk over a critical or sensitive period of the life course, a MR estimate should capture the causal effect of this biomarker but may not be able to distinguish period effects | Negative exposure control design |
| Weak instrument bias | If IV is not robustly associated with the exposure, estimates will be biased towards the observational estimate in a one-sample setting and towards the null in a two-sample setting | Increase sample size<br>Genetic risk scores<br>Two-sample MR analysis |
| “Winner’s Curse” | Chance correlation between genetic variants and confounders can introduces an overestimation of the effect of a “lead” genetic variant on an exposure of interest in the discovery stage of a GWAS. The effect of this phenomenon will depend on the degree of overlap of participants in the GWAS discovery dataset and subsequent MR analyses. In a one-sample MR setting with a binary outcome, winner’s curse should not lead to bias if control participants were used in the discovery GWAS. If both cases and controls were used in the discovery dataset, this will lead to weak instrument bias. If the instrument is identified in a sample independent to the one in which MR analysis is performed, this will lead to an underestimate of the causal effect. | Two-sample MR analysis<br>Split-sample MR analysis |
| Low statistical power | Genetic variants typically explain a small amount of variance for a given exposure, thus MR requires large sample sizes to test hypotheses with adequate power. Furthermore, in finite samples, confounders may not be perfectly balanced between genotypic groups | Large GWAS and GWAS consortia<br>Genetic risk scores<br>Two-sample MR analysis |

### Comparison of Mendelian randomization to Randomized Controlled Trials

Due to the random allocation of alleles at conception it can be useful to compare the structure of a MR analysis to the design of a randomized trial, where individuals are randomly allocated at baseline to an intervention or control group (**Figure 1**). Groups defined by genotype should be comparable in all respects (e.g., approximately equal distribution of potential confounding factors) except for the exposure of interest. It follows that any observed differences in outcomes between these genotypic groups can be attributed to differences in long-term exposure to the trait of interest. This latter point is an important distinction when interpreting results from a MR analysis as compared to a randomized controlled trial: MR will generally estimate the effect of life-long “allocation” to an exposure on an outcome, unless an exposure typically occurs only from a certain age - e.g., alcohol consumption and smoking - and the genetic proxy affects metabolism of that exposure ^34^. If the effect of this exposure on an outcome is cumulative over time, a MR analysis may generate a larger effect estimate than that which would be obtained from a randomized trial examining an intervention over a limited duration of time. Additionally, if the effect of an exposure on an outcome operates primarily or exclusively over a critical or sensitive period of the life course (e.g., early childhood), a MR analysis should be able to “capture” a causal effect of this exposure but will not be able to distinguish such period effects. In contrast, a randomized trial will have the flexibility to test certain interventions over restricted periods of follow-up and in individuals who may be within narrow age ranges. These distinctions are discussed in more detail in “Cancer Latency and Reverse Causation – benefits of MR”.

**Figure 1.**
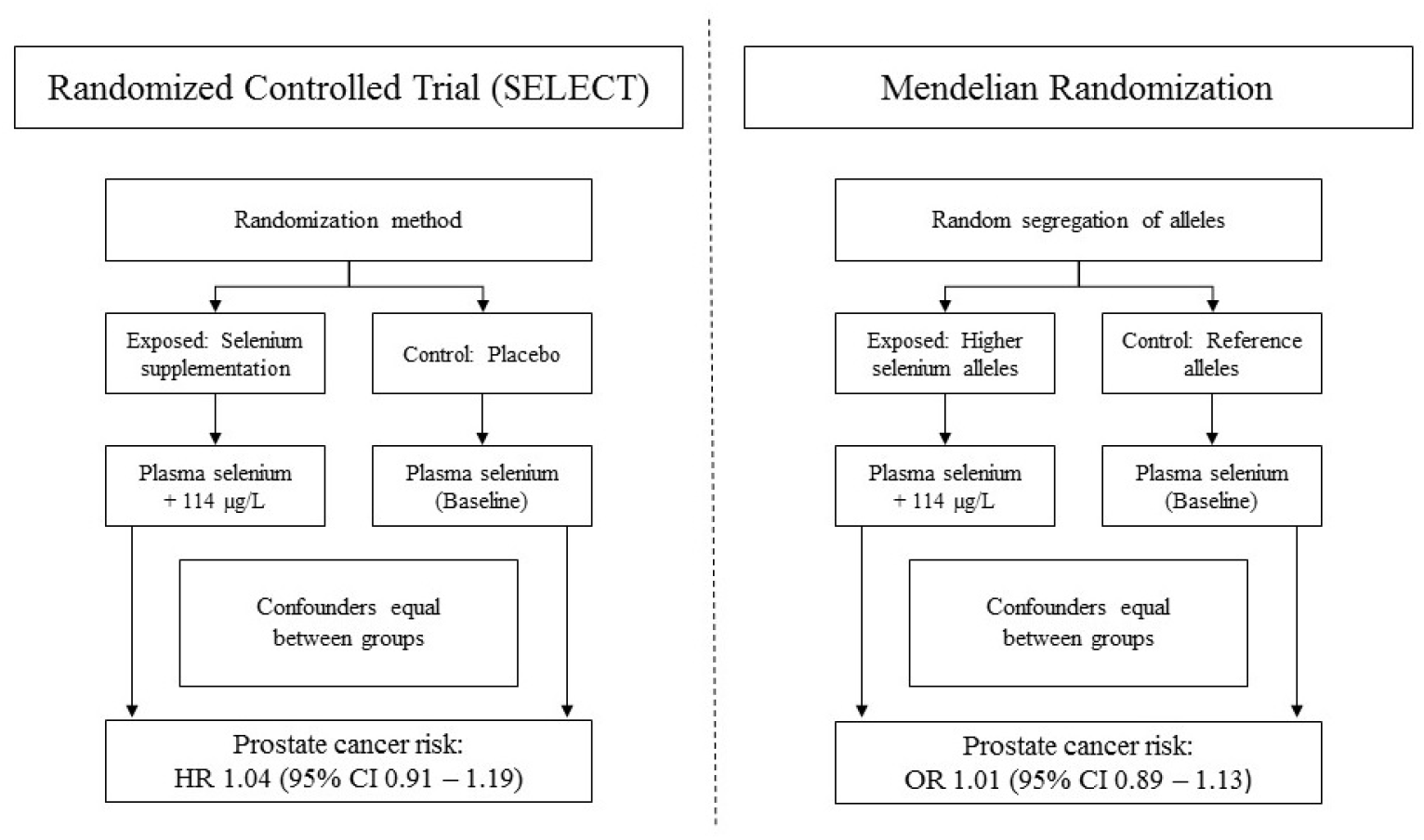
Schematic comparison of the structure of a randomized controlled trial (SELECT) and a Mendelian randomization analysis (PRACTICAL) In SELECT (left), individuals were randomly allocated to the intervention (200 μg daily selenium supplementation, which lead to a 114μg/L increase in blood selenium) or control group (placebo). In PRACTICAL (right), the additive effects of selenium-raising alleles at eleven SNPs, randomly allocated at conception, were scaled to mirror a 114μg/L increase in blood selenium. If an RCT trial is adequately sized, randomization should ensure that intervention and control groups are comparable in all respects (e.g., distribution of potential confounding factors) except for the intervention being tested. In an intention-to-treat analysis, any observed differences in outcomes between intervention and control groups can then be attributed to the trial arm to which they were allocated. Likewise, in a MR analysis, groups defined by genotype should be comparable in all respects (e.g., distribution of both genetic and environmental confounding factors) except for their exposure to a trait of interest. Any observed differences in outcomes between groups defined by genotype can then be attributed to differences in life-long exposure to the trait of interest under study.

More formally, MR is a form of instrumental variable (IV) analysis that relies on three key assumptions: the IV (here, one or more genetic variants) should (i) be reliably associated with the exposure of interest; (ii) not be associated with any confounding factor(s) that would otherwise distort the association between the exposure and outcome; and (iii) should not be independently associated with the outcome, except through the exposure of interest (known as the “exclusion restriction criterion”) (**Figure 2a**). If all assumptions are met, MR can provide an unbiased causal estimate of the effect of an exposure on disease or a health-related outcome. Violation of one or more of these assumptions means that instruments are invalid and, consequently, that findings from such an analysis may yield a biased effect estimate.

**Figure 2.**
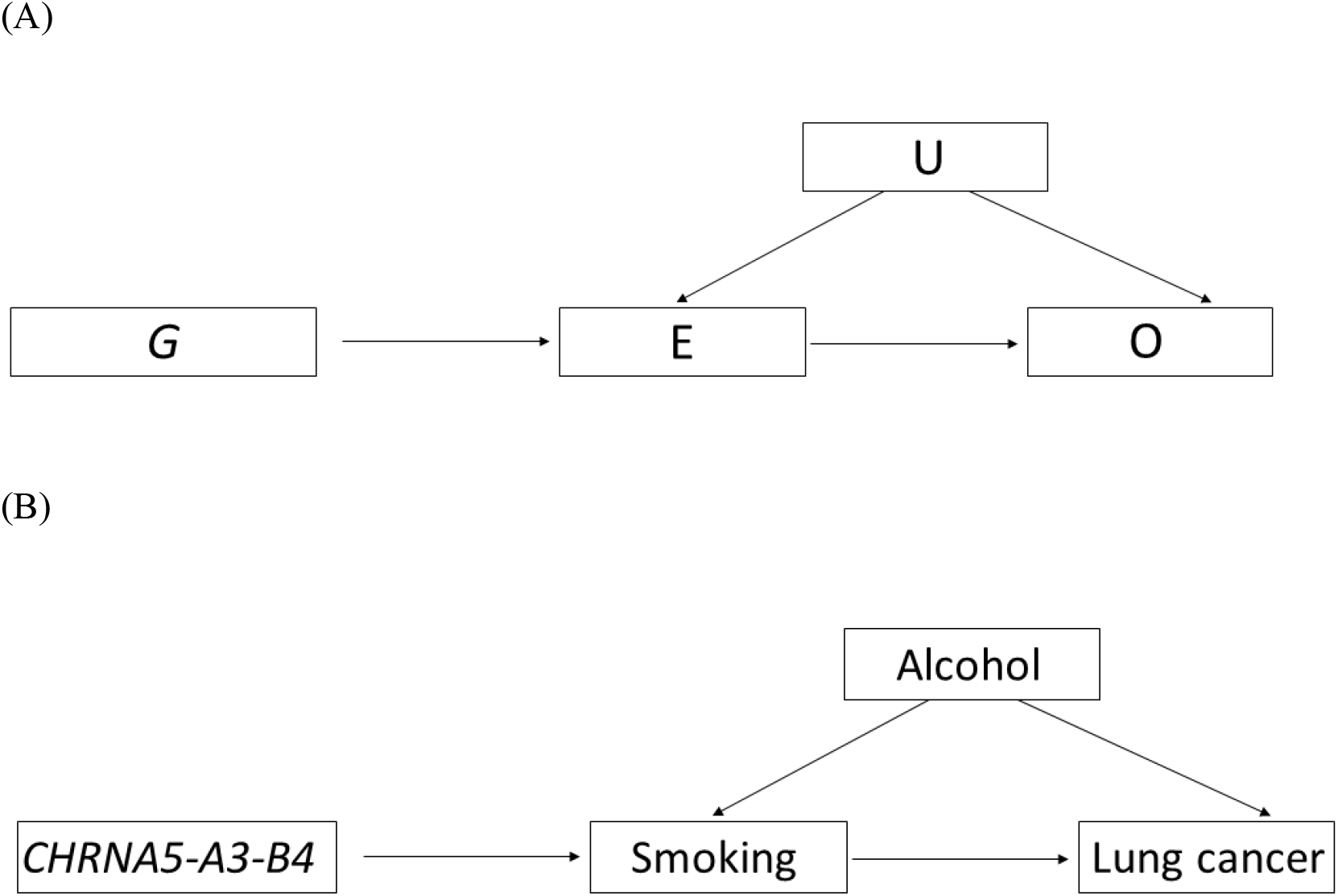
Illustration of MR methodology. (A) A genetic variant (G) is used as a proxy for a modifiable exposure (E) to assess the association between E and an outcome of interest (O) without the issues of reverse causation, and confounding (U). MR methodology relies on three main assumptions, in that G must (i) be reliably associated with E; (ii) not be associated with U; and (iii) not be independently associated with O, except through E. This method is exemplified in the context of assessing the association of smoking and lung cancer (B), using the *CHRNA5-A3-B4* SNP as a genetic instrument for heaviness of smoking.

### Previous success of Mendelian randomization approaches and potential for cancer research

Over the past decade, MR has been increasingly adopted as an analytical approach within population health research, particularly the fields of metabolic and cardiovascular disease (CVD), where there are several notable examples of important causal inferences. For example, MR has suggested a likely causal role of statins on type 2 diabetes (T2D) risk ^35,36^; likely non-causal roles of circulating levels of high-density lipoprotein cholesterol (HDL-C) in CVD ^37^ and C-reactive protein (CRP) in T2D ^38^; pointed to the efficacy of proprotein convertase subtilisin/kexin type 2 (PCSK9) inhibitors for CHD prevention prior to the publication of confirmatory long-term trial results ^35,39^; and prioritized further examination of apolipoprotein B ^40,41^, lipoprotein(a) ^42^ and interleukin-6 ^43^ and de-prioritized fibrinogen ^44^ and secretory phospholipase A(2)-IIA ^45^ as intervention targets for CVD. Although this approach has scope to test the causal effects of an increasing number of exposures relevant to cancer through the continued growth in large-scale GWAS output (**Box 1**), to date there remains a noticeable gap in the MR literature with regard to cancer compared to other outcomes (**Box 2**).

#### Box 1: Instrumentable exposures

Types of cancer-relevant exposures with robust genetic associations, which therefore could be instrumented in a MR context include: i) behavioural and lifestyle exposures (e.g., alcohol consumption, nutrient biomarkers, milk and caffeine consumption, lifecourse sun exposure); ii) endogenous biomarkers (e.g., fatty acids, glycaemic traits, insulin, interleukin-6, insulin-like growth factor, CRP, sex-steroid hormones, vitamin D, adiponectin); iii) drug targets (e.g., 3-hydroxy-3-methylglytaryl-CoA reductase (HMGCR), prostaglandin endoperoxidase synthase 2 (PTGS2), proprotein convertase subtilisin/kexin type 9 (PCSK9)); iv) site-specific DNA methylation markers; v) small metabolites (e.g., amino acids, lipids, lipoproteins, steroids); and vi) early life factors (e.g., birthweight, childhood obesity, and age at puberty onset).

#### Box 2: Existing publications applying Mendelian randomization within a cancer context

The number of MR studies published annually has increased rapidly since the early 2000s, reaching approximately 226 publications in 2016 alone. However, the proportion of these studies examining the causal effects of one or more traits on cancer incidence or progression has remained modest (see **Figure** below).

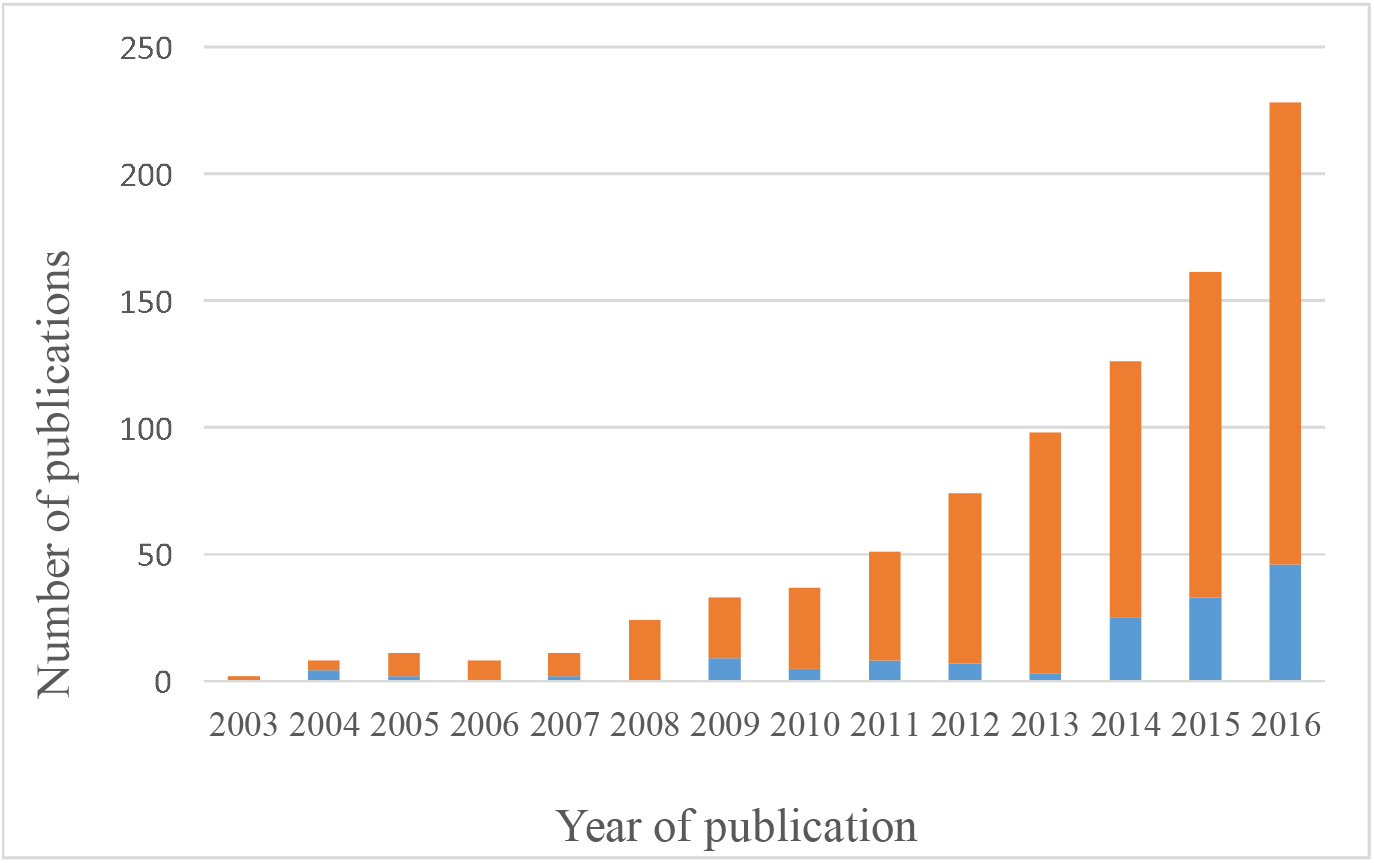

Using results from PubMed, the figure above represents the proportion of all published MR studies per year that assessed cancer incidence or progression as an outcome. Blue bars represent the number of all MR cancer studies (all MR studies=blue bars + orange bars) published from 2003-2016. PubMed search strategy for all MR studies: (mendelian randomization analysis[MeSH Terms]) OR “mendelian randomization” OR “mendelian randomisation”; PubMed search strategy for MR cancer studies: (((cancer) OR neoplasms[MeSH Terms])) AND (((mendelian randomization analysis[MeSH Terms]) OR “mendelian randomization”) OR “mendelian randomisation”)

Here, we provide an overview of some recent studies that have applied MR to cancer outcomes, highlighting both the potential strengths compared to conventional epidemiological studies and the unique challenges of performing MR studies in cancer. Recent methodological extensions to the original MR paradigm are presented, with emphasis on the translational opportunities that they may offer to inform drug target validation and public health strategies to reduce the burden of cancer.

### Considerations for MR in cancer

Both the principal strengths of MR and important limitations of this method have been discussed in detail previously ^26–32,46–50^. The latter are presented in **Table 1** with some methodological and statistical approaches that have been developed to address them outlined in **Box 3** and **Box 4**. Some considerations which are specific to investigating causality in the setting of cancer are outlined below.

#### Box 3: Summarized data and two-sample MR

##### Two-sample MR

Historically, both gene-exposure and gene-outcome estimates in MR analyses had to be obtained from a single sample which relied upon the availability of information on genotype, exposure, and outcome among all participants in that dataset. In practice, this not only posed a challenge in that large-scale measurement of a given exposure of interest (e.g., many molecular traits) may not only be prohibitively expensive but also that measurement of certain exposures may not be possible (e.g., if adequate blood sample collection or preservation has not taken place) ^124^. An extension to the original MR paradigm that has allowed MR analyses to overcome some of these challenges is the integration of gene-exposure and gene-outcome estimates from two independent (non-overlapping) datasets into a single analysis, an approach called “two-sample MR” analysis ^124,125^.

##### Two-sample MR with summarized genetic association data

It is possible and increasingly common practice to perform MR analyses exclusively using summarized data on gene-exposure and gene-outcome estimates ^125,126^. A strength of two-sample MR with summary data is that the scope of possible MR analysis can be expanded significantly by exploiting the growing amount of publicly-available summary data from large genome-wide association study (GWAS) consortia ^127^ and is aided by the development of a harmonised MR platform that has collated these datasets (MR-Base) ^128^. Utilizing data from separate exposure and outcome samples can help to bolster statistical power in MR analyses by increasing the overall sample size of an analysis, particularly when testing effects on binary disease outcomes like cancer, and also reduces the likelihood of “winner’s curse” bias (see Table 1) ^125^. This increased power also means that sensitivity analyses to test pleiotropy assumptions (see Box 4) which are often statistically inefficient are better-powered to detect violations of these assumptions. Furthermore, whereas in a one-sample MR setting weak instruments can bias effect estimates towards the observational effect, resulting in potential false positive associations, in a two-sample setting weak instrument bias distorts findings towards the null. Thus, conducting both analyses is a form of sensitivity analysis that provides bounds to a possible causal effect.

To test whether height has a causal effect on risk of colorectal, lung, and prostate cancer, Khankari *et al*. used a two-sample MR approach. This employed: i) summarized gene-exposure estimates from a panel of 423 single-nucleotide polymorphisms (SNPs) previously found to be associated with height in a large GWAS meta-analysis (GIANT consortium; N=253,288) and collectively explaining approximately 16% of variance in height; and ii) summarized gene-outcome estimates from a total of 47,800 cancer cases (across the three outcomes ascertained) and 81,533 controls from the Genetic Associations and Mechanisms in Oncology (GAME-ON) consortium ^129^. This approach allowed robust causal inference with adequate statistical power. While Khankari *et al*. did not examine the effects of height across stage/grade or histological sub-type of the three cancers examined, two-sample approaches enable statistically efficient examination of risk factors across such stratified groups which may have limited sample sizes.

##### Limitations of two-sample MR

While two-sample MR offers some clear advantages over a conventional one-sample approach, it also introduces additional assumptions. One important assumption is that the separate datasets from which gene-exposure and gene-outcome associations are obtained are representative of the same underlying population, for example with regard to sex, age, ethnicity, or genetic profile. While most GWAS that have examined sex-specific associations of traits have often reported at most modest evidence of sexual dimorphism ^130,131^, given the sex-specific nature of certain cancers, care should be taken to ensure that instruments are obtained from sex-stratified GWAS for analyses of these cancers when available. For example, in examining the effect of waist-hip-ratio (WHR) on endometrial or ovarian cancer this could involve using the 34 SNPs associated with WHR in women exclusively as a primary instrument, then comparing results with those obtained using the 47 SNPs associated with WHR across both sexes as a sensitivity analysis ^132,133^. Concordance of findings between both approaches may suggest that directionally-consistent SNPs associated with WHR at genome-significance in women, but not men, simply reflected reduced statistical power in sex-stratified GWAS analyses and not genuine heterogeneity in SNP-effects between sexes. A second challenge when performing two-sample MR using summary data is the difficulty in examining the IV assumption that an instrument used is independent of exposure-outcome confounders. While restriction of analyses to ethnically homogenous gene-exposure and gene-outcome datasets will reduce the possibility of confounding through population stratification, in lieu of data on measured potential confounders, this assumption cannot be directly tested. While one way of approximately testing this assumption is performing look-up of associations of SNPs with suspected potential confounders in curated GWAS databases, this would not preclude chance confounding relationships arising in the dataset(s) from which summary data were obtained. Third, with the use of summary data from large GWAS consortia, it is possible that there may be some participant overlap in the datasets from which gene-exposure and gene-outcome associations are obtained. If overlap is small, this should not substantially bias effect estimates, however substantial overlap will bias MR toward the observational effect ^134^.

#### Box 4: Genetic risk scores and pleiotropy

##### Using multiple genetic variants as an instrument

While GWAS over the past decade have been successful at identifying robust associations between common genetic variants (usually SNPs) and thousands of phenotypes, the effects of individual variants on traits are often modest ^135^. Consequently, statistical power for MR analyses using single variants as instruments can be limited. A common approach of overcoming limited statistical power is to combine multiple variants into a genetic risk score (GRS), which increases the variance explained for a trait of interest, improving instrument strength ^136,137^. A GRS can consist of an unweighted summation of risk-factor increasing alleles across variants but, more commonly, a weighted approach is used (e.g., weighted by the inverse of the standard error of the gene-outcome association – called the “inverse-variance weighted (IVW) method”). In a two-sample setting (see Box 3), a GRS will typically be constructed by combining SNPs that are independent (i.e., not in LD with each other) into a weighted score. However, it is also possible to combine correlated SNPs in low to moderate LD into a GRS, using weighted generalized linear regression for example ^136^. This requires the creation of a weighting matrix which takes into account correlations between SNPs, often with use of a reference panel like the Hapmap or the 1,000 Genomes Project ^138,139^, which is then used to correctly inflate standard error estimates. The latter method may be preferable to overcome weak instrument issues when few independent SNPs are available.

##### Vertical vs horizontal pleiotropy

While construction of a GRS can help to enhance statistical power in MR analyses, increasing the number of variants included in a score is accompanied by an increased probability that any of these variants could be pleiotropic (i.e., one variant having effects on two or more traits). In a genetic epidemiological context, an important distinction is made between vertical and horizontal pleiotropy, each having different effects on the interpretation of MR findings. Vertical pleiotropy occurs when one variant has an effect on two or more traits that both influence an outcome through the same biological pathway. For example, variants in *FTO* that not only associate with BMI, but also with fasting insulin and glucose concentrations would be consistent with a causal effect of BMI on these downstream traits ^140^. In this case, a MR analysis examining the effect of BMI on T2D risk using these *FTO* variants would be consistent with an instrument (genetic variants associated with BMI) influencing an outcome (T2D) exclusively through the exposure of interest (BMI). This form of pleiotropy would be expected in complex biological systems and does not pose a threat to the validity of a MR analysis ^141^. In contrast, horizontal pleiotropy occurs when one variant has an effect on two or more traits that influence an outcome through independent biological pathways. For example, genetic variants associated with triglyceride levels also show substantial overlap with variants associated with LDL-C and HDL-C ^142^. As a putative effect of triglyceride-increasing variants on CHD risk may not only operate through elevation of triglycerides but through alternate cholesterol pathways, a naïve MR analysis using all triglyceride-increasing variants without addressing pleiotropy in this instance could invalidate the “exclusion restriction criterion” IV assumption. The presence of horizontal pleiotropy thus poses a direct threat to the validity of MR findings.

##### Assessment of horizontal pleiotropy

When using either a single or a small number of genetic variants as IVs, the presence of horizontal pleiotropy for any individual variant can be assessed through SNP look-ups in curated GWAS databases with complete summary data (e.g., MR-Base ^128^, PhenoScanner ^143^, dbGap ^144^) to examine whether associations for a given SNP have been reported for traits other than the exposure of interest. Sensitivity analyses can then be performed by dropping variants that are suspected to be horizontally pleiotropic and then carefully interpreting pooled causal estimates with and without suspected horizontally pleiotropic SNPs. When an instrument consists of multiple genetic variants, an important first step in examining the presence of horizontal pleiotropy in analyses is to assess heterogeneity in causal estimates across individual IVs (including visually examining heterogeneity using a funnel plot). While substantial heterogeneity in causal estimates may be indicative of the presence of horizontal pleiotropy, if there is overall symmetry in the funnel plot, pleiotropic effects will be balanced (termed “balanced pleiotropy”) and the overall causal estimate generate will be unbiased. In contrast, if there is considerable asymmetry in a funnel plot, this will suggest that horizontal pleiotropic effects of individual IVs are not balanced and that overall causal estimates will be biased (termed “directional pleiotropy”). MR-Egger regression and the weighted median estimator (WME) are two widely implemented approaches for detecting and accounting for directional pleiotropy, and are applicable to analyses utilizing individual-level and summary-level data ^121,122^. An additional approach called the mode-based estimate (MBE) has also recently been proposed as a method to examine horizontal pleiotropy in MR analyses ^145^. All of these methods can help to detect IV violations while making different assumptions about the nature of horizontal pleiotropy and thus, when feasible, using all approaches as sensitivity analyses in a given MR analysis can serve as an important mechanism to assess the robustness of findings to pleiotropic bias.

##### Sensitivity analyses to examine horizontal pleiotropy when using multiple genetic variants

MR-Egger regression provides a consistent causal effect estimate even when all genetic variants are invalid IVs because they violate the exclusion restriction criterion. This approach performs a weighted linear regression of the gene-outcome coefficients on the gene-exposure coefficients with an unconstrained intercept term. If the IV assumption that the association of each variant with the outcome is mediated exclusively through the exposure of interest is met, this intercept term should be zero. An intercept term that differs from zero would suggest the presence of unbalanced pleiotropy, thus providing a test for directional pleiotropy. In turn, the slope coefficient in MR-Egger regression will provide an estimate of a causal effect adjusted for directional pleiotropy. An important consideration when using MR-Egger is that it works under the InSIDE (instrument strength independent of direct effect) assumption. In essence, InSIDE assumes that no association exists between the strength of gene-exposure associations and the strength of bias due to horizontal pleiotropy. Intuitively, if multiple genetic variants in an MR analysis have horizontally pleiotropic effects through unrelated intermediate variables, it would be expected that this assumption should hold ^7^. However, this assumption is unlikely to be satisfied in situations where all pleiotropic effects are due to the presence of a single confounder. As such, in lieu of an established method of formally testing the InSIDE assumption, interpretation of intercept terms and slope coefficients generated through MR-Egger should be made with this assumption in mind. A complementary sensitivity analysis to MR-Egger is the weighted median estimator. This approach provides an estimate of the weighted median of a distribution in which individual IV causal estimates in a risk score are ordered and weighted by the inverse of their variance. Unlike MR-Egger which can provide an unbiased causal effect even when all IVs are invalid, WME requires that at least 50% of the information in a risk score is coming from IVs that are valid in order to provide a consistent estimate of a causal effect in a MR analysis. However, an advantage of WME is that it provides improved precision as compared to MR-Egger and does not rely on the InSIDE assumption. The mode based estimator generates a causal effect using the mode of a smoothed empirical density function of individual IV causal estimates in a risk score. This approach operates under the assumption that the most common effect estimate of individual IVs in a risk score arises from valid instruments (called the Zero Modal Pleiotropy Assumption, or ZEMPA). If this assumption holds, the mode can provide a consistent causal estimate even if most of the (non-modal) IVs are invalid. Both simple and weighted mode approaches (weighted by the inverse variance of the SNP-outcome association) can be utilized. Mode-based approaches have less power to detect a causal effect than the weighted median estimator but greater power than MR-Egger regression under the condition of no invalid instruments. Similar to the weighted median estimator, mode-based approaches are also (by default) less susceptible to bias from outlying variants in a risk score.

### Cancer Latency and Reverse Causation – benefits of MR

Given long latency periods for many cancers, spurious findings resulting from reverse causation (when the direction of cause-and-effect relationship is contrary to the presumed direction) are an important concern in cancer epidemiology. Reverse causation has been suspected in several instances of ambiguous ^51–53^ or paradoxical findings ^54^ in the cancer literature. For example, early studies documenting an association between higher circulating cholesterol and lower cancer incidence were variably interpreted as plausible evidence of a protective effect of raised cholesterol on cancer risk or as latent cancer leading to a reduction in cholesterol levels ^55–57^ With the introduction and widespread usage of LDL-C lowering medications for the prevention and treatment of CVD, concern arose that such measures could thus be increasing cancer rates ^58,59^.

In an early proposal of the use of genetics as a tool to circumvent issues of reverse causation in observational data, Katan *et al*. ^60^ suggested examining the association of genetic variants in the *APOE* locus, determinants of circulating cholesterol levels, with cancer risk. As germline genotype at *APOE* was fixed at conception, it was argued that it would not be influenced by subsequent cancer development and could therefore be used to establish whether cholesterol had a causal effect on cancer incidence (**Figure 3a**). Subsequent MR analyses testing the effect of lifelong elevated cholesterol through genetic variation in *APOE, NPC1L1, PCSK9*, and *ABCG8* have reported null associations with overall cancer risk ^61–63^. These findings alongside secondary analyses of statin trials showing no effect on cancer rates ^64^ suggest that – a potential explanatory role of confounding aside - early observational findings supporting a protective effect of cholesterol on cancer risk likely reflected undiagnosed cancer or early carcinogenic processes causing a reduction in cholesterol levels in pre-diagnostic samples.

**Figure 3.**
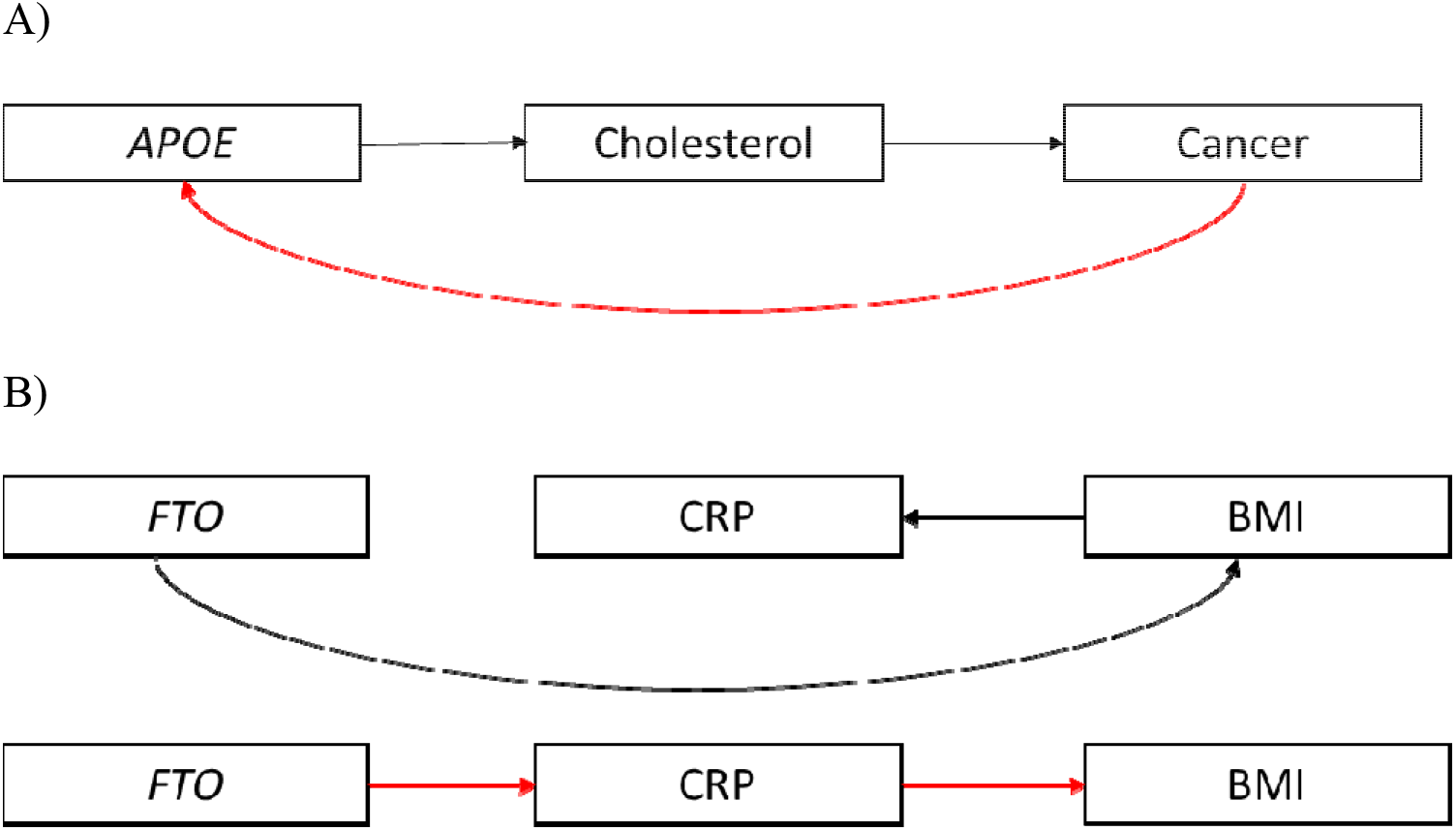
Reverse causation within Mendelian randomization studies. A) SNPs within *APOE* are used as instruments for cholesterol to assess the causal effect of cholesterol on cancer risk in a MR framework. *APOE* is a determinant of differential plasma cholesterol levels and is fixed at conception; therefore, will not be influenced by cancer (red arrow). B) A SNP which primarily influences BMI (e.g. *FTO*) will also influence CRP due to the causal effect of BMI on CRP. However, if this SNP is mistakenly used as an instrument for CRP then this will lead to erroneous results (red arrow).

### Long-term exposure – benefits of MR

The advantages of exploiting the fixed nature of germline genotype extends beyond addressing reverse causation in observational studies. Large cancer prevention trials are often constrained to examining interventions over a limited duration in time and over a particular period in the life-course (e.g., middle and/or late adulthood) ^65^. Given the length of time required for solid tumor development ^66^, randomized trials will often not allow sufficient follow-up for the effect of an intervention to be detected. In turn, long-term chemoprevention trials that are conducted may suffer from issues of non-compliance in the intervention arm, contamination in the control arm, and attrition during follow-up.

Further, the optimal timing of an exposure to prevent cancer may be early in the life-course and therefore may not be adequately addressed in randomized trials ^67^. For example, it has been proposed that certain carcinogenic agents or processes may confer an effect, or a particularly pronounced effect, only over ‘critical periods’ of early life or adolescence (e.g., the influence of inadequate childhood nutrient intake on adult cancer risk or the pubertal period as a window of breast cancer susceptibility) ^68–72^. Interrogating the long-term effect on cancer of a given intervention in a prevention trial among children or adolescents would be unfeasible.

Examining the effect of genetic variants allocated at conception can therefore offer an important first step in identifying risk factors that may be sensitive to duration or timing of an exposure over the life course. Inferences made from promising MR findings to plausible intervention effects in a subsequent randomized trial would then need to carefully consider the possibility that effect estimates obtained in a MR analysis could be sensitive to critical period effects (in which case intervening on an exposure outside of this period may not alter disease risk) or represent the cumulative effect of lifelong exposure to a biomarker (in which case a relatively short-term trial may generate a smaller effect estimate than that obtained from MR). Adopting a “triangulation” framework where evidence from different epidemiological approaches with non-overlapping sources of bias are integrated can then be used to further examine durations of intervention necessary to confer an effect or ‘pinpoint’ possible critical or sensitive windows of susceptibility to carcinogenic agents ^73^. For example, multivariable regression analyses examining the association of an exposure, with some evidence of causality from MR studies, over different lengths of follow-up may help to identify the duration of exposure required to confer an effect. In contrast, a negative control study with repeat measures of an exposure both within and outside of hypothesized critical or sensitive periods (e.g., dietary fat intake before, during, and after pubertal development), in relation to subsequent disease risk (e.g., breast cancer)^74^ can help refine periods of increased vulnerability to cancer-causing exposures.

### Cancer Latency and Reverse Causation – limitations of MR

Genetic variants known to directly affect an exposure will in some cases be well-characterized (e.g., variants in the *APOE* locus), and it will be established whether or not the variant-exposure associations are influenced by the outcome of interest. The biological understanding of other variants associated with risk factors that are identified in GWAS, however, is often more limited. In some situations in which genetic variants are associated with both a proposed exposure and outcome of interest, the association between genetic variant and outcome might be via the exposure (i.e., a valid IV analysis) but it is also possible that, under certain circumstances, there may be a primary effect of the genetic variant on the outcome which in turn causes a change in the exposure. This can potentially bias MR estimates in both one-sample and two-sample analyses.

This situation has been illustrated previously in the context of body mass index (BMI) and C-reactive protein (CRP) where an erroneous causal effect can be generated if a genetic variant that primarily influences BMI, which in turn influences CRP levels because BMI has a causal effect on CRP, is mistaken as being a variant with a primary influence on CRP (**Figure 3b**). ^26^ Use of such a variant as an instrument for CRP in a MR analysis of the effect of CRP on BMI would then lead to biased results.

This introduction of reverse causation into a MR analysis may be problematic for common cancers with long latency periods between tumour initiation and diagnosis (e.g., breast and prostate) ^75^. Reverse causation in this context could be mitigated by obtaining gene-exposure estimates in a healthy population where the prevalence of undiagnosed, latent cancer is likely to be low. These estimates could then be used to generate IV estimates in a two-sample MR framework. Additionally, steps could be taken to construct a GRS solely consisting of instruments that plausibly act directly on a trait. For example, in constructing an instrument for CRP levels, this could include solely using variants within the *CRP* gene itself as these variants are more likely to be exclusively associated with CRP levels than variants in other genes ^76^. However, it should be noted that a trade-off of using few, biologically-informed SNPs as an instrument is that sensitivity analyses examining horizontal pleiotropy (e.g., MR-Egger, median-based methods, mode-based methods) – when feasible to perform (i.e., at minimum, the availability of three or more SNPs) – will have limited statistical power to detect a causal effect.

### Selection bias in cancer progression analyses

A particular concern in cancer epidemiology is that exposures that influence cancer incidence may not influence cancer progression or survival. For example, although smoking is a robust risk factor for breast cancer incidence, smoking cessation upon development of breast cancer seems to have little effect on subsequent survival ^77^. There has been some suggestion that folate may play a dual role in prostate and colorectal carcinogenesis: protective against DNA damage prior to the development of neoplasia, but promoting tumour progression via enhanced tumour proliferation and tissue invasion once cancer has developed ^78,79^.

Some MR studies have begun to examine the effect of risk factors on both cancer incidence and progression ^80^. In a recent analysis examining the effect of alcohol consumption on prostate cancer risk in 46,919 men in the PRACTICAL consortium, alcohol consumption (instrumented by 68 SNPs in alcohol-metabolizing genes) was reported not to be associated with overall prostate cancer risk but to confer an increased risk of prostate cancer mortality among men with low-grade disease ^81^. Such MR studies exploit the fact that GWAS are being increasingly used to identify genetic variants associated with cancer progression or survival ^82,83^. While these studies have to date generally uncovered few genome-wide significant loci, GWAS association estimates can still be useful for identifying causal risk factors for progression (particularly with use of a two-sample MR framework).

However, there are important methodological considerations in investigating factors causing cancer progression. This is because prognostic studies can suffer from selection bias due to the fact that any factors that cause disease incidence (or diagnosis) will tend to be correlated with each other in a sample of only cases, even when they are not correlated in the source population. Thus if at least one factor causes both incidence and disease survival (hypothetically, insulin resistance in **Figure 4**), all the other factors which cause disease incidence (hypothetically, smoking in **Figure 4**) will appear to be associated with survival, unless the true prognostic factor is conditioned upon. Thus, the estimated effect on progression for any factor that is associated with incidence is likely to be biased. However, any factor that is not associated with incidence will not suffer from selection bias by studying only cases in a MR analysis. For example, given evidence for a causal effect of BMI on breast cancer incidence ^84^, in their MR analysis of BMI on breast cancer survival Guo et al. had to consider the possibility that evidence of an effect of BMI on breast cancer survival could reflect confounding with one or more other causes of breast cancer incidence that became conditionally associated with BMI upon restricting analyses to breast cancer cases ^85^.

**Figure 4.**
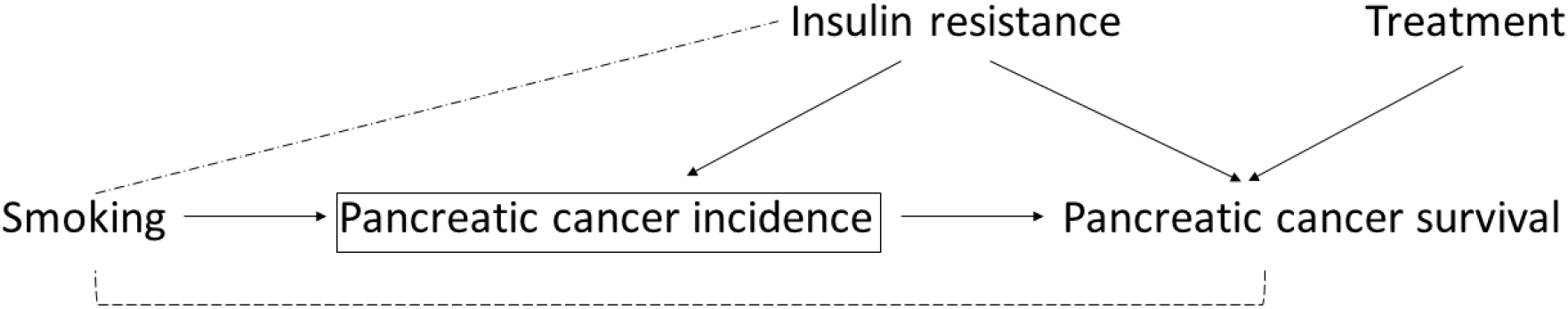
Directed acyclic graph for selection bias in prognostic studies. In this example, the square bracket indicates that we are conditioning on pancreatic cancer incidence in a survival study by only studying pancreatic cancer cases, thus inducing an association between smoking (a factor that is otherwise independent of pancreatic cancer survival) and pancreatic cancer survival. This link is broken when conditioning on the factor that influences both cancer incidence and survival (e.g., insulin resistance), which can otherwise be seen as a confounder of the association between smoking and cancer survival. If a factor appears to influence pancreatic cancer survival that is not associated with pancreatic cancer incidence (e.g., treatment for pancreatic cancer), selection bias in such an MR analysis would not be expected.

When conducting prognostic studies, care should be taken to examine and (where possible) overcome the selection bias due to studying only cases ^80^. First, the observed data could also be used to help identify plausible directed acyclic graphs (DAGs) including both disease incidence and progression. For example, if a genetic score for a phenotype, and an environmental variable, are correlated in cases, but not in the source population this would suggest that both factors influence disease incidence, diagnosis, or self-selection into the study. However, lack of evidence for such correlations does not imply that there is no selection bias, and expert or external knowledge should be used in constructing the DAG, as is usual practice. The DAG can then be used to help inform sensitivity analyses. Additional data on factors that predict incidence could be combined with observed data in cases, to minimise selection bias, either by conditioning or by inverse probability weighting (IPW). If more than one DAG are considered plausible a priori, then they can be used to conduct sensitivity analyses, by examining how robust the conclusions are to the causal assumptions made. The DAG can also be used to identify which assumptions are being made that are untestable given the observed data, and then sensitivity analyses can be conducted by examining plausible values for those relationships.

### Illustrative examples

To illustrate the use of MR in analyses examining cancer outcomes, we have outlined three studies that have employed this approach to understand the causal role of various exposures on cancer incidence.

#### Selenium and incidence of prostate cancer

Prospective studies reporting inverse associations of dietary, blood, and toenail selenium with risk of prostate cancer ^86–92^, along with findings from *in vitro* studies ^93,94^, led to development of the Selenium and Vitamin E Cancer Prevention Trial (SELECT) ^95^. SELECT was a 2 × 2 factorial trial of 35,533 healthy middle-aged men that examined the effect of daily supplementation with selenium, vitamin E, or both agents combined, as an intervention for prostate cancer prevention. The trial was stopped after 5.5 of a planned 12 years follow-up due to a lack of efficacy compounded by possible carcinogenic (increased rates of high-grade prostate cancer) and adverse metabolic (weak increased rates of T2D) effects in the selenium supplementation group ^9,10^. It is plausible that residual confounding may have accounted for conflicting results between prospective studies and SELECT ^96,97^.

To test whether a MR approach could have predicted the results of SELECT, a two-sample MR analysis (Box 3) was performed using summary data on 72,729 individuals of European descent from the PRACTICAL consortium ^98,99^ Eleven SNPs robustly associated with blood selenium in a meta-analysis of previously published GWAS ^100,101^ (*P* < 5 × 10^−8^) were combined into a GRS (**Box 4**) to proxy circulating levels of selenium (**Figure 1**). To allow for direct comparison of effect estimates with SELECT, the authors investigated the causal odds ratio (OR) per 114 μg/L genetically-elevated circulating selenium, scaled to match the measured differences in blood selenium between supplementation and control arms in SELECT.

Consistent with results from SELECT, a 114 μg/L life-long increase in blood selenium in MR analyses was not associated with overall prostate cancer risk (OR: 1.01, 95% CI 0.89–1.13; *P* = 0.93; SELECT: Hazard Ratio (HR): 1.04, 95% CI 0.91–1.19). MR analysis of selenium on advanced prostate cancer (defined as Gleason ≥8, prostate-specific antigen >100 ng/mL, metastatic disease (M1), or death from prostate cancer) (OR: 1.21, 95% CI 0.98–1.49; *P* = 0.07) was concordant with the observed weak evidence for an increased risk of high-grade prostate cancer (Gleason ≥7) in the selenium supplementation arm of SELECT (HR: 1.21, 95% CI 0.97–1.52; *P* = 0.20). Likewise, the effect of selenium on T2D (OR: 1.18, 95% CI 0.97–1.43; *P* = 0.11) was concordant with weak evidence for an increased risk of T2D in the selenium arm of SELECT (HR: 1.07, 95% CI 0.97-1.18; *P* = 0.16).

Thus, the overall similarities in findings between this MR analysis and that of SELECT, as compared to results from conventional observational studies, provides some support for the utility of an MR approach in approximating experimental results using observational data. Further, these results suggest that performing a MR analysis may be an important time-efficient and inexpensive step in predicting both efficacy and possible adverse effects of an intervention before an RCT is performed. This information, along with careful consideration of the possibility that an MR result could be driven by critical period effects (e.g., limiting benefit of intervention in middle-aged adults) or represent the cumulative effect of long-term exposure to a biomarker (i.e., limiting benefit of a trial with limited duration), could be used to help prioritize which interventions should be taken forward to a trial.

#### Alcohol and incidence of oesophageal cancer

Regular alcohol consumption is associated with a substantial increased risk of oesophageal squamous cell carcinoma in observational studies, with an approximate two-fold increased risk for moderate drinkers and a five-fold increased risk for heavy drinkers when compared to occasional/non-drinkers ^102^. However, alcohol consumption is often associated with other lifestyle and behavioural factors (e.g., smoking and dietary intake), which may themselves predispose toward oesophageal cancer ^103,104^. Further, most studies that examined this hypothesis have used case-control designs, which may introduce reporting bias if cases recall alcohol consumption differently from controls. For example, cases may be more likely to reflect on and more carefully report their history of alcohol consumption, to account for their cancer diagnosis, than controls who do not have the same motivation for more careful recall of alcohol exposure ^102^. Given the high global prevalence of alcohol consumption, considerable population-level reductions in the incidence of oesophageal cancer could be achieved by intervening on alcohol consumption levels if the causal nature of the link between the two was confirmed ^105^.

The ability to metabolize acetaldehyde, the principal metabolite of alcohol and a carcinogen ^106^, is encoded by *ALDH2*, which is polymorphic in some East Asian populations. Specifically, the *ALDH2* *2 allele produces an inactive protein subunit that is unable to metabolize acetaldehyde, resulting in markedly higher peak blood alcohol levels in *2*2 homozygotes compared to *1*1 homozygotes ^107^. Individuals with the *2*2 genotype experience a flushing reaction to alcohol, along with dysphoria, nausea, and tachycardia, and therefore have very low levels of alcohol consumption ^108^. Consequently, genetic variation in *ALDH2* can be utilized as an instrument for examining both regular alcohol consumption and blood acetaldehyde levels among alcohol consumers ^109^.

In a meta-analysis of seven studies with a total of 905 oesophageal cancer cases of East Asian descent, individuals with the *ALDH2* *2*2 genotype were found to have an approximately 3-fold reduced risk of oesophageal cancer, as compared to the *ALDH2* *1*1 genotype (OR: 0.36, 95% CI 0.16–0.80), suggesting a protective effect of reduced alcohol consumption on oesophageal cancer risk ^110^. However, when comparing individuals with a heterozygous *1*2 genotype to *1*1 individuals, the former were shown to have a (seemingly paradoxical) overall increased oesophageal cancer risk (OR: 3.19, 95% CI 1.86–5.47). A naïve interpretation of this finding, without consideration of the effect of the *ALDH2* *2 allele on blood acetaldehyde, would suggest that individuals with moderate alcohol intake had the highest risk of oesophageal cancer.

When this association was stratified by self-reported alcohol intake, the effect of *1*2 genotype on oesophageal cancer was shown to differ markedly by alcohol intake. Among non-drinkers, there was no strong evidence for an increase in risk among heterozygotes (OR: 1.31, 95% CI 0.70–2.47) relative to *1*1 individuals. However, among heavy drinkers there was a 7-fold increase in risk (OR: 7.07, 95% CI 3.67–13.6). Similarly, meta-regression analysis showed evidence that level of alcohol intake influenced the effect of the *1*2 genotype on oesophageal cancer risk (*P* = 0.008) (i.e., the larger the amount of alcohol intake, the greater the OR of *1*2 versus *1*1 genotypes). As the possession of an *ALDH2* *2 allele only appeared to increase risk of oesophageal cancer among heterozygotes who reported alcohol intake, this suggested that the substantially elevated acetaldehyde levels in these heterozygotes may mediate the effect of alcohol intake on oesophageal cancer. Subsequent MR analysis examining the effect of alcohol intake on risk of head and neck cancer similarly reported both protection of *ALDH2* *2*2 genotype against head and neck cancer as compared to *1*1 homozygotes and an interaction of the effect of *ALDH2* 1*2 genotype on cancer risk by alcohol behaviour (elevated risks among moderate and heavy drinkers compared to *1*1 individuals, but no elevated risk among non-drinkers) ^111^.

More generally, this example illustrates how interpretation of MR findings can be challenging when there is limited biological understanding of the genetic variant used as a proxy for a given exposure. MR results that appear to be strongly discordant with underlying biology should be followed-up alongside available functional understanding of genetic variants employed as instruments to help resolve ambiguous or paradoxical results and avoid naïve interpretation of findings.

#### Body mass index and incidence of lung cancer

In contrast to the relationship of adiposity with risk of most cancers, BMI has shown consistent inverse associations with incidence of lung cancer, particularly among current and former smokers ^112,113^. As smoking is a robust risk factor for lung cancer and has been shown to have an inverse effect on BMI ^114^, some have argued that residual confounding by smoking could account for this apparent protective association ^115^. Reverse causation (i.e., undiagnosed lung cancer or disease processes leading up to lung cancer prior to study entry influencing subsequent weight loss), especially in cohorts with insufficient follow-up time, has also been proposed as an explanation for this observational finding ^116^.

Attempts to address these possible sources of bias have failed to provide clarity. For example, studies that reported finely stratifying associations across various dimensions and classifications of smoking behaviour (e.g., number of cigarettes smoked per day, “cigarette-years” smoked, and time since quitting smoking) have found little evidence to support residual confounding by smoking influencing this association ^112,113^. Further, studies removing individuals with inadequate follow-up have reported little effect on overall findings ^112,113,117,118^, interpreted as suggesting that reverse causation is unlikely to be a major contributor to this association. For this reason, some have argued that the inverse association of BMI with lung cancer may not be artefactual but may reflect a real biological phenomenon ^113^.

Given that germline genetic variants associated with BMI cannot be influenced by prevalent disease and should not be associated with potential confounding factors, a MR approach could be used to assess whether increased BMI is protective against lung cancer ^84,119^. For example, Carreras-Torres *et al*. performed a MR analysis using GWAS results on 16,572 lung cancer cases and 21,480 controls of European descent ^120^. 97 SNPs previously associated with BMI in a GWAS of 339,224 individuals from the Genetic Investigation of Anthropometric Traits consortium were compiled into a GRS to proxy for anthropometrically measured BMI. This risk score was associated with measured BMI but not with available measures of tobacco exposure, including pack-years, cigarettes smoked per day, or cotinine levels, providing some evidence against confounding through measured smoking variables ^114^. In two-sample MR analyses, a 1-SD increase in BMI was weakly associated with an increased risk of lung cancer (OR: 1.13, 95% CI 0.98–1.30; *P* =0.10), with strong heterogeneity across histological sub-types (*P*_heterogeneity_ < 3 × 10^−5^). Notably, BMI was positively associated with risk of both squamous cell (OR: 1.45, 95% CI 1.16–1.62; *P* = 1.2 × 10^−3^) and small cell carcinoma (OR: 1.81, 95% CI 1.14–2.88; *P* = 0.01) but showed weak evidence for a protective effect for adenocarcinoma (OR: 0.82, 95% CI 0.66–1.01; *P* = 0.06). These findings were robust to sensitivity analyses (MR-Egger regression and WME) ^121,122^ performed to identify potential directional pleiotropic effects of the GRS (see Box 4). These findings thus help to clarify a likely positive risk relationship of genetically-elevated BMI with two major histosubtypes of lung cancer. Alongside some genetic evidence to suggest that elevated BMI may influence subsequent smoking uptake ^123^, which itself reduces BMI while increasing lung cancer risk ^114^, these findings collectively suggest a possible mechanism that could help to reconcile seemingly conflicting MR and observational findings. Further interrogation of a possible mediating role of smoking on the causal pathway between BMI and lung cancer risk using “two-step MR” (discussed in "MR for mediation") may be able to help shed further light on the possible intricate relationship between smoking and BMI in the aetiology of lung cancer.

### Recent methodological extensions and future applications

In recent years, the development of various methodological extensions to the original MR paradigm have helped to enhance the scope of MR analyses, several of which are discussed below with reference to possible applications in cancer epidemiology.

### MR for mediation

Over the past decade, high through-put “omics” technologies have begun to permit exhaustive profiling of the epigenome, metabolome, and proteome (as examples), allowing the collection of high-dimensional molecular data on increasingly large number of individuals ^146^. Such omics measures may serve as important mediators on causal pathways linking macro-level risk factors with cancer incidence or progression. Identification of these omics mediators could help to increase scope for pharmacological intervention aimed at cancer prevention or progression (e.g., when upstream risk factors are difficult or not feasible to intervene on). While conventional mediation analyses exist to examine possible exposure-mediator-outcome relationships, the validity of these approaches relies upon strong assumptions which are unlikely to be met in practice, such as no measurement error and no unmeasured confounding ^147^.

With the performance of GWAS on large collections of metabolites and other omic measures ^148,149^, this will create opportunity to develop genetic instruments for these traits. To establish whether a particular molecular intermediate is on the causal pathway between an exposure and cancer, genetic variants can be used as instruments for both exposures and putative mediators that influence a disease outcome in a two-step MR framework (**Figure 5**) ^150^.

**Figure 5.**
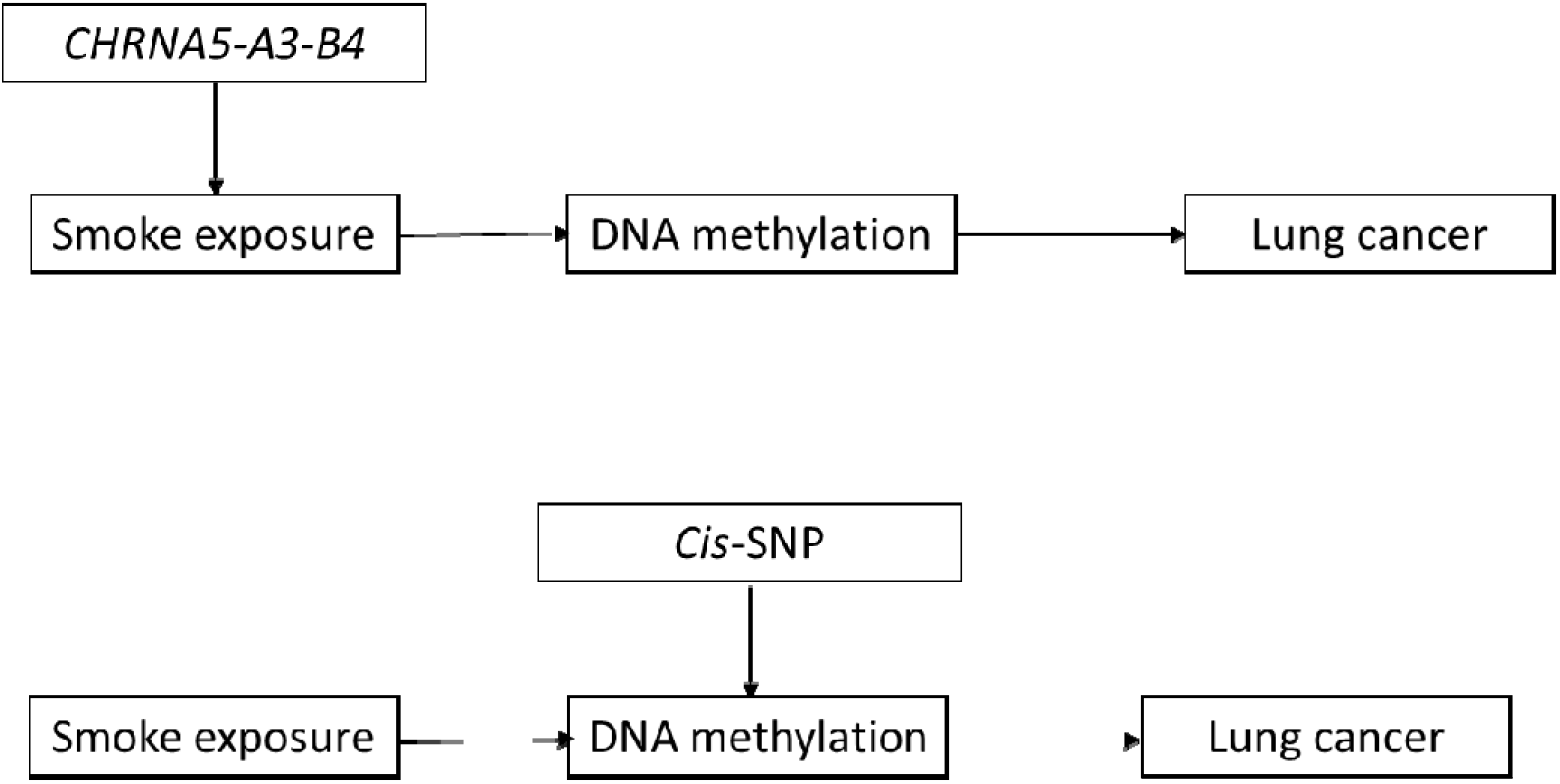
Two-step Mendelian randomization analysis examining the mediating effect of methylation on the association between smoke exposure and lung cancer. In the first step, a SNP within *CHRNA5-A3-B4* is used as an instrument for smoke exposure to assess the causal association between smoking and DNA methylation. In the second step, an independent cis-SNP is used as an instrument for DNA methylation to assess the causal association of DNA methylation with lung cancer risk. The two-step method allows interrogation of the mediation effect of DNA methylation in the association between smoking and lung cancer risk.

For example, a method of testing the mediating role of methylation changes on cancer outcomes would be to exploit the fact that genetic variants (e.g., methylation quantitative trait loci, mQTLs) are robustly associated with methylation at CpG sites across the epigenome, providing possible instruments for MR analyses ^151^. Two-step MR could then used to examine the potential mediating role of DNA methylation sites associated with exposures such as tobacco smoke ^152^ which have also been found to be strongly associated with lung cancer risk ^153^. To test whether methylation is causally mediating (some, or all of) the effect of tobacco exposure on lung cancer risk, in the first step, a SNP could be used to proxy smoking behaviour in order to investigate its effect on the intermediate phenotype (DNA methylation). In the second step, an independent SNP (not related to the exposure) could then be used to proxy the intermediate phenotype (DNA methylation) which could then be examined in relation to the disease outcome (lung cancer) ^147^.

### ExE interaction

Akin to a factorial RCT, factorial MR is a method of testing the independent and additive effects of two or more exposures (ExE) on disease outcomes (**Figure 6**). This approach was adopted by Ference *et al*. who performed a 2×2 factorial MR analysis to examine the effect of the LDL cholesterol-lowering drug ezetimibe on risk of CHD, as compared to the effect of statins alone or when combined with statins ^154^. Ference *et al*. examined the effect of natural random allocation to lower LDL-C on the risk of CHD through SNPs in *NPC1L1* (a target of ezetimibe) alone, *HMGCR* (a target of statins) alone, or variants in both gene regions combined. The authors reported that natural randomization to lower LDL-C through SNPs in *NPC1L1* and *HMGCR* alone showed similar decreases in LDL-C and CHD and that randomization to lower LDL-C in both groups combined had a linearly additive effect on LDL-C lowering and a log-linearly additive effect on CHD risk. These results were corroborated by the ‘Improved Reduction of Outcomes: Vytorin Efficacy International Trial,’ which allocated 18,144 participants to ezetimibe, statins, both, or placebo ^155^.

**Figure 6.**
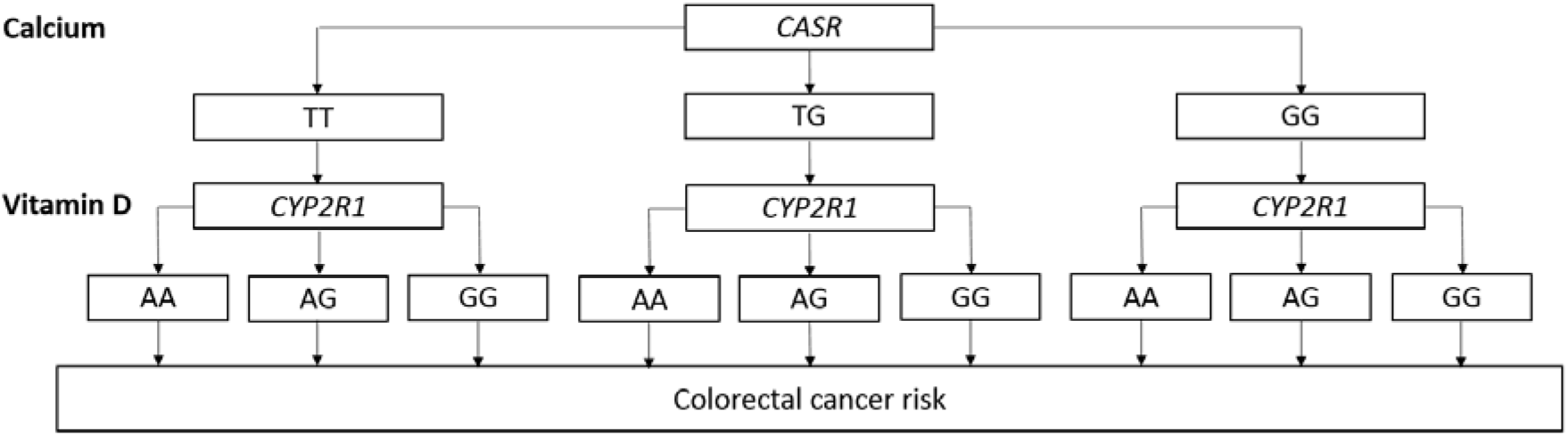
Factorial Mendelian randomization applied to the effects of calcium and vitamin D on colorectal cancer risk. This hypothetical example of a factorial MR trial uses genetic proxies for calcium (e.g., variants in *CASR*) and vitamin D (e.g., variants in *CYP2R1*) to examine the possible independent and additive effects of these exposures on colorectal cancer risk.

Factorial MR could be used to evaluate the proposed interactions between smoking and alcohol consumption on risk of head and neck cancer ^156^, smoking and BMI on risk of oesophageal cancer ^112^, or vitamin D and calcium on risk of colorectal cancer ^157^, as examples. An important caveat of this approach is that it relies on access to individual-level data and requires very large sample sizes to have adequate statistical power to reliably detect differences in effect across groups.

It may also be useful (particularly for exposures that cannot be instrumented) to embed MR analyses into randomized trials as another method of simulating a factorial randomized trial. For example, data from the Add-Aspirin trial, which is examining whether regular aspirin use after curative treatment for early stage solid tumours can prevent disease recurrence and survival, could present an opportune setting to examine possible additive effects of aspirin and other therapies (proxied by genetic instruments) on cancer outcomes ^158^. Such an embedded MR-in-RCT design would necessitate large sample sizes and would require access to individual-level trial data.

### GxE interaction

Studies of gene-environment interactions (GxE) offer the potential to discover both novel genetic risk loci and environmental factors for cancer, in addition to gaining insight into biological mechanisms that may underlie associations between risk factors and cancer ^159^. In turn, these findings can be used to inform risk prediction and the identification of high-risk populations for targeted prevention strategies ^160^. As in conventional observational studies, the “E” component of GxE studies will still be subject to the same methodological limitations that undermine robust causal inference. The use of genetic variants to instrument the “E” component in these studies can therefore help to circumvent confounding and bias in conventional observational analyses ^161^. Limited statistical power is an important consideration in conventional GxE studies and the reduced power that would accompany the use of genetically-instrumented exposures should also be considered.

GxE interaction can also be used to test violations of pleiotropy assumptions in certain MR analyses. Specifically, as MR models assume that a genetic instrument is not associated with an outcome except through modulation of the exposure of interest, this assumption can be tested by stratifying analyses on a supposed interacting variable. For example, in a MR analysis of the effect of heaviness of smoking on lung cancer risk, an instrument proxying for heaviness of smoking (variants in *CHRNA5-A3-B4*) ^162^ should only be associated with lung cancer among self-reported smokers and not in never or former smokers in an analysis stratified by smoking status ^30^. Likewise, in a MR analysis of the effect of alcohol on oesophageal or head and neck cancer, an instrument proxying for regular alcohol consumptions (variants in *ALDH2*) should only be associated with these cancers among those who report regularly consuming alcohol and not among those who refrain from consuming alcohol ^110,111^. Here, the interaction reflects the lack of any pathway through which the genotype can operate if the exposure is not present (i.e. a lack horizontal of pleiotropy) and allows for the exclusion restriction criterion to be tested. When using GxE to test pleiotropy assumptions in this manner, however, it is important to consider the possibility of introducing selection bias (see “Selection bias in cancer progression analyses”).

### Hypothesis-free MR

A novel extension to a conventional “hypothesis-driven” MR analysis is a phenome-wide, “hypothesis-free” MR analysis (termed “MR-PhEWAS”) ^163^. This approach makes use of genotyped datasets with high-dimensional phenotypic data or summary GWAS association statistics to perform hundreds or thousands of statistical tests simultaneously in an agnostic manner. For example, the approach can be used to examine the causal effect of a single exposure across multiple outcomes or multiple exposures across a single outcome. In contrast to hypothesis-driven analyses, hypothesis-free approaches allow for testing hypotheses that may not have been considered or tested previously, thus identifying novel risk relationships, and can help to address issues of publication bias as all analyses are openly specified and all results are presented) ^164^.

For example, using a two-sample MR framework with summary data, Haycock *et al*. performed a MR-PheWAS examining the causal effect of telomere length on risk of 35 cancers and 48 non-cancer diseases in 420,081 cases and 1,093,105 controls ^165^. After correction for multiple-testing, they found that telomere length increased cancer risk across most sites and histological sub-types but reduced CVD risk. An important consideration when performing hypothesis-free MR analyses using summary data is the need to follow-up any putative findings in subsequent independent datasets. This can be a challenge when using summary GWAS data to perform such analyses if a large proportion of the available GWAS literature was used to provide causal estimates in the original “discovery phase” of an analysis.

### MR for identifying causality of mutational signatures

Large-scale analysis of the genomes of thousands of cancer patients has helped to reveal somatic “mutational signatures” (distinctive somatic mutational patterns left by unique carcinogenic agents) involved in the development of their tumours ^166,167^. To date, mutational signatures have been identified across more than 30 different cancer types, with anywhere from two to six distinction mutational processes for each cancer type. Knowledge of the causes of somatic mutations within tumour tissue can improve understanding of the mechanisms by which endogenous and exogenous exposures promote the development of a cancer. Of the mutational signatures identified across cancer types, a putative cause has been proposed for approximately half ^166^; MR may offer particular promise in helping to identify the aetiology of other mutational signatures identified ^168^.

Robles-Espinoza *et al*. examined the effect of germline *MC1R* status, associated with red hair, freckling, and sun sensitivity, on somatic mutation burden in melanoma. Such an analysis can be viewed as a MR appraisal of the effect of this sensitivity phenotype on somatic mutation burden in melanoma ^169^. For all six mutational types assessed, there was evidence of an increased burden of somatic single nucleotide variants in individuals carrying one or two *MC1R* R alleles (disruptive variants). For one of the six mutational signatures characterized by an abundance of somatic C>T single nucleotide variants, each additional R allele at *MC1R* was associated with a 42% (95% CI 15-76%) increase in the C>T single nucleotide variant count. This approach therefore highlights the possibility of testing the causal effect of suspected carcinogenic agents on mutational burden for various mutational signatures across cancer tissues and sub-types.

### Drug repurposing and adverse drug effects

Drug repurposing, applying known drugs to novel indications, can provide a rapid, cost-effective mechanism for drug discovery and may hold promise for the development of pharmacological interventions for cancer prevention ^170,171^. In turn, for well-tolerated drugs that are considered candidates for repurposing, MR may offer an attractive approach for testing their potential chemopreventive efficacy. For example, it is currently possible to reliably instrument drugs for which there is a broad understanding of the biological mechanism of action: examples that currently exist within cardiovascular disease include proxies for HMG Co-A reductase inhibitors, PCSK9 inhibitors, CETP inhibitors, and sPLA2 inhibitors ^172^. For the primary or tertiary prevention of certain cancers, aspirin, metformin, and bisphosphonates have all been proposed as possible candidate pharmaceutical agents for repurposing ^173–175^. Using MR as a first step to test drug efficacy for novel cancer indications could help to prioritize or deprioritize further investigation of certain drugs (e.g., helping to guide which drugs should be taken forward to testing in RCTs for re-purposing).

MR may also provide a useful approach for predicting adverse effects of pharmaceuticals ^176^. Pre-approval trials are often not able to adequately capture development of adverse effects due to the comparatively small number of individuals typically exposed to a drug in such trials (unless drug effects are very common or very large), the limited duration of most trials, the possibility that recorded data may not include necessary information to identify unanticipated drug effects or those unrelated to the drugs’ indication, and unknown generalizability of trial participants to the broader population that they are meant to represent. While many of these issues can be addressed post-approval of a drug through spontaneous reporting systems, these introduce their own limitations including confounding, for example by indication, environmental factors, or lifestyle traits. MR studies should be able to overcome these limitations and have been employed in some instances to test or anticipate adverse effects of interventions in ongoing intervention trials ^35,177^. For example, MR studies examining genetic variants in *HMGCR* as an instrument for statin exposure have corroborated findings from large phase III trials that statins modestly increase risk of type 2 diabetes ^36,178,179^.

While knowledge of biological pathways can help to anticipate some adverse drug effects pre-approval of a drug, it may not be possible to correctly predict all such effects ^180^. One possible approach to resolve this would be to use MR-PhEWAS to perform a phenotypic scan of a genetically-instrumented drug exposure across hundreds or thousands of potential outcomes, as outlined previously. The identification of possible adverse effects of a drug through this approach could then be used to pre-specify and adequately power secondary outcome measures or, alternately, to de-prioritize further investigation of a therapeutic target. For drugs currently on the market, MR can be used to test for potential adverse effects that were not picked up during the trial, in addition to appraising causality of effects that have been reported through conventional reporting systems.

### Conclusion

Observational epidemiological studies are prone to various intractable biases which can undermine robust causal inference. Mendelian randomization offers a promising approach to generate a more reliable evidence-base for cancer prevention and treatment. The advent of MR methods using summarized data means that such analyses can now be performed more efficiently, rapidly, and with greater statistical power than previously possible. Further, the range of methodological extensions to the original MR paradigm now available have greatly expanded the scope of this approach, enabling increasingly sophisticated causal questions to be interrogated ^181^. Given optimism surrounding use of the method in helping to strengthen evidence for public health and pharmacological interventions ^182^, it is likely that there will be a continued proliferation of MR analyses in the literature in the near future. Careful design, analysis, and interpretation of such studies with consideration of the limitations of the method will provide the greatest opportunity for such studies to inform cancer prevention and treatment strategies.

### Funding

The MRC IEU is supported by the Medical Research Council and the University of Bristol (MC_UU_12013/1-9). The Integrative Cancer Epidemiology Programme is supported by Cancer Research UK programme grant C18281/A19169. J.Y. and R.L. are funded by CRUK PhD studentship C18281/A20988. C.J.B. is funded by the Wellcome Trust 4-year studentship WT083431MA

